# Prophage-dependent recombination drives genome structural variation and phenotypic heterogeneity in *Escherichia coli* O157:H7

**DOI:** 10.1101/2020.12.02.407981

**Authors:** Stephen F. Fitzgerald, Nadejda Lupolova, Sharif Shaaban, Timothy J. Dallman, David Greig, Lesley Allison, Sue C. Tongue, Judith Evans, Madeleine K. Henry, Tom N. McNeilly, James L. Bono, David L. Gally

## Abstract

The human zoonotic pathogen *Escherichia coli* O157 is defined by its extensive prophage repertoire including those that encode Shiga toxin, the factor responsible for inducing life-threatening pathology in humans. As well as introducing genes that can contribute to the virulence of a strain, prophage can enable the generation of large-chromosomal rearrangements (LCRs) by homologous recombination. This work examines the types and frequencies of LCRs across the major lineages of the O157 serogroup and defines the phenotypic consequences of specific structural variants. We demonstrate that LCRs are a major source of genomic variation across all lineages of *E. coli* O157 and by using both optical mapping and ONT long-read sequencing demonstrate that LCRs are generated in laboratory cultures started from a single colony and particular variants are selected during animal colonisation. LCRs are biased towards the terminus region of the genome and are bounded by specific prophages that share large regions of sequence homology associated with the recombinational activity. RNA transcriptional profiling and phenotyping of specific structural variants indicated that important virulence phenotypes such as Shiga toxin production, type 3 secretion and motility are affected by LCRs. In summary, *E. coli* O157 has acquired multiple prophage regions over time that act as genome engineers to continually produce structural variants of the genome. This structural variation is a form of epigenetic regulation that generates sub-population phenotypic heterogeneity with important implications for bacterial adaptation and survival.

**Author Summary:** *Escherichia coli* has an ‘open genome’ and has acquired genetic information over evolutionary time, often in the form of bacteriophages that integrate into the bacterial genome (prophages). *E. coli* O157 is a clonal serogroup that is found primarily in ruminants such as cattle but can cause life-threatening infections in humans. *E. coli* O157 isolates contain multiple prophages including those that encode Shiga-like toxins which are responsible for the more serious disease associated with human infections. We show in this study that many of these prophages exhibit large regions of sequence similarity that allow rearrangements to occur in the genome generating structural variants. These occur routinely during bacterial culture in the laboratory and the variants are detected during animal colonization. The variants generated can give the bacteria altered phenotypes, such as increased motility or toxin production which can be selected in specific environments and therefore represent a highly dynamic mechanism to generate variation in bacterial populations without a change in overall gene content.

## Introduction

Bacterial viruses, termed prophage, that incorporate their genomes onto the bacterial chromosome are major drivers of bacterial genome evolution, host and niche adaptation and virulence [1–3]. Prophage integration directly benefits the bacterial host by conferring resistance against other lytic viruses [4], by carriage of virulence factors, including toxins and effector proteins [1, 5], enzymes involved in stress resistance [6] and the expression both gene regulators and sRNAs capable of influencing the host gene regulatory network [2, 7]. Here we examine the impact prophages have on the structure of the bacterial genome through the generation of large-chromosomal rearrangements (LCRs).

*Escherichia coli* O157:H7 is a significant human zoonotic pathogen originating from ruminant hosts, especially cattle [8]. Over evolutionary time, numerous prophage (typically 16 – 25) have integrated into the genomes of *E. coli* O157 strains with an integration bias towards the terminus (Ter) of replication [9]. Acquisition of these prophage, many of which are closely related λ-like phage, has driven the evolution of this pathogen by carriage of virulence genes including secreted effector proteins, sRNAs involved in virulence gene regulation and [7, 10], importantly, these prophage include those that encode Shiga toxin (Stx) subtypes. Stx toxins are the main mediators of vascular endothelial cell killing in infected humans [11] and the resulting damage can lead to haemolytic uremic syndrome (HUS), often fatal, or lead to life-long kidney and brain damage [12–14]. *E. coli* O157:H7 strains are divided into three phylogenetically distinct lineages (I, I/II and II) with those that represent a serious threat to human health belonging to lineage I or Lineage I/II and the majority encode two sub-types of Stx, Stx2a and Stx2c. Stx2a is generally associated with more serious disease [11, 15–18] and the emergence of *E. coli* O157:H7 as a zoonotic threat correlates with the introduction of Stx2a-encoding prophage into the *E. coli* O157 cattle population approximately 50 years ago [11].

There is published evidence that *E. coli* O157 type strain EDL933 can undergo large-chromosomal rearrangements (LCRs), mainly inversions [19, 20], with these rearrangements being flanked by prophages. LCRs, such as inversions, duplications and translocations, occur by homologous recombination between repeat sequences on the same chromosome [21]. While LCRs arising between ribosomal *rrn* operons, pathogenicity islands and insertion sequence (IS) elements have been associated with speciation, diversification, outbreaks and immune evasion in bacteria [1, 22] few studies have examined LCRs arising from inter-prophage recombination and their impact on phenotype.

In this study we demonstrate that prophage-mediated LCRs are a major source of genomic variation across all lineages of *Escherichia coli* O157. We show that alternate chromosomal conformations are generated during laboratory culture and are selected during host colonisation. Specific LCRs were associated with changes in virulence phenotypes and we therefore propose that the generation of LCRs within *E. coli* O157 populations *in vivo* facilitates phenotypic heterogeneity and niche adaptation, include host colonisation. Prophage act as genome engineers by driving conservative rearrangements leading to sub-populations with distinct phenotypes that can provide an advantage in different environments.

## Results

### LCRs shape *E. coli* O157 genome evolution

To examine the extent of genomic diversity generated by LCRs in the *E. coli* O157 clonal group, we examined the whole genome sequences of 72 isolates, the majority of which were generated by PacBio long-read sequencing (Table S1). Strains analysed were representative of the main *E. coli* O157 lineages (I, I/II and II) and included multiple sub-Lineage Ic, PT21/28 isolates which have been responsible for the majority of serious human infections in the UK over the last two decades [11]. This genome dataset included previously sequenced complete genomes from each lineage, including strains Sakai (NC_002695.2), EDL933 (CP008957.1) and TW14359 (CP001368) (Table S1).

Pairwise alignment of all 72 genomes identified LCRs, predominantly large inversions, as a common source of genomic variation between isolates within each *E. coli* O157 lineage with the exception of lineage I/II (Supplementary Figure S1). In addition, each genome was individually aligned against a representative reference strain from each of four lineages and the chromosomal loci of all LCRs >50 kb were mapped (Fig 1A-D). The reference strains were: Strain 9000 (Lineage 1c), Sakai (Lineage 1a), TW14359 (Lineage I/II) and Strain 180 (Lineage II).

**Figure 1.**
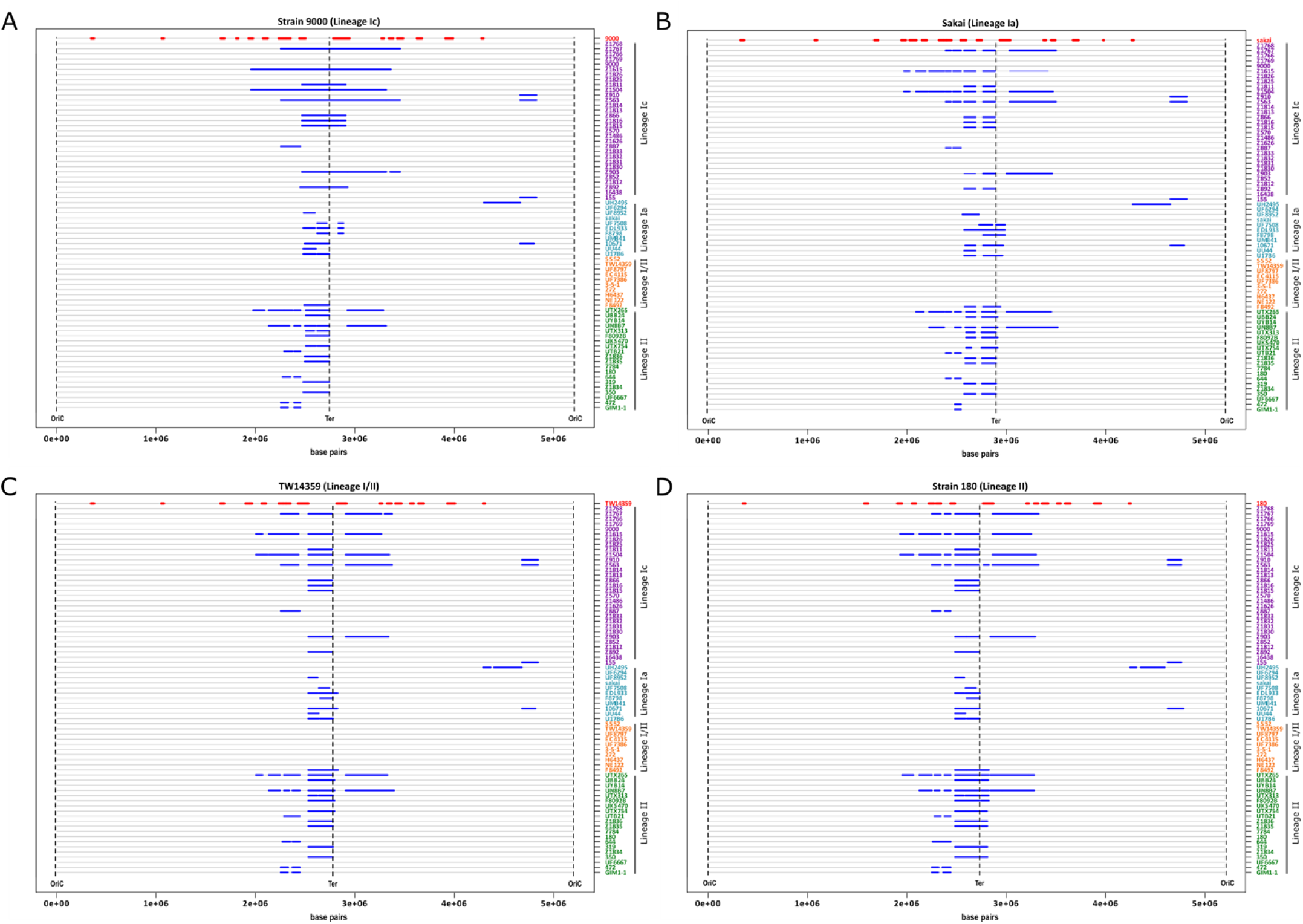
Position of major chromosomal rearrangements in *E. coli* O157 genomes. The relative positions of all LCRs ≥ 50 kb (blue lines) are marked on the chromosomal maps (grey line) of strains from Lineage 1c (purple), Lineage 1a (blue) Lineage I/II (orange) and Lineage II (green). Chromosomes are centred by the replication terminus (Ter), beginning and ending at the origin of replication (OriC). LCRs are shown relative to four reference strains: (**A**) 9000, Lineage 1c; (**B**) Sakai, Lineage Ia; (**C**) TW14359, Lineage I/II; (**D**) 180, Lineage II. The position of the main prophage (red line) are mapped for each comparison strain.

LCRs > 50 kb were frequently identified in lineages Ia, Ic and II irrespective of the reference strain used for alignment, however it was evident that Lineage I/II strains exhibited less variation (Figure 1). Strains from Lineage 1c and Lineage II exhibited the most variation at this macro level with an average of 43 and 37 LCRs identified, respectively, (Table. 1 and Fig. 1). Strains from Lineage 1a were less variable with an average of 14 LCRs identified across all strains and the least genomic variation with respect to the reference strains was observed for strains from Lineage I/II with an average of just 2.5 LCRs identified in a single strain, F8492. We note that strain F8492 was a singleton isolate that grouped closely with our other representative Lineage I/II strains (Supplementary Figure S2). Lineage I/II strains were also the least variable when the number of LCRs identified were corrected to account for the unequal number of strains analysed within each lineage (Table. 1). To examine this further, we plotted the average size of all LCRs with a lower cut-off of >20 kb that could be detected in each strain relative to the four reference genomes (Supplementary Figure S2 A – D). At this lower cut-off, LCRs ranging between 20 kb and 30 kb were identified in Lineage I/II that were generally consistent across all Lineage I/II strains relative to each reference genome. These results indicate that the macro genome conformation of Lineage I/II strains is highly conserved. While LCRs can occur within Lineage I/II strains they have a reduced capacity to generate larger LCRs > 30 kb compared with the two other lineages.

**Table 1.** Mean number and size of LCRs relative to each reference genome

| Reference | Total No. of LCRs |  |  |  |
| --- | --- | --- | --- | --- |
|  | Lineage Ia | Lineage Ic | Lineage I/II | Lineage II |
| 9000 | 18 | 36 | 2 | 43 |
| Sakai | 15 | 47 | 2 | 38 |
| TW14359 | 13 | 47 | 2 | 34 |
| 180 | 13 | 43 | 4 | 41 |
| Mean | 14.75 | 43.25 | 2.5 | 39 |
| Reference | LCRs per lineage/strain |  |  |  |
|  | Lineage Ia | Lineage Ic | Lineage I/II | Lineage II |
| 9000 | 1.64 | 1.16 | 0.2 | 2.15 |
| Sakai | 1.36 | 1.52 | 0.2 | 1.9 |
| TW14359 | 1.18 | 1.52 | 0.2 | 1.7 |
| 180 | 1.18 | 1.39 | 0.4 | 2.05 |
| Mean | 1.34 | 1.40 | 0.25 | 1.95 |
| Reference | Average LCR length (bp) |  |  |  |
|  | Lineage Ia | Lineage Ic | Lineage I/II | Lineage II |
| 9000 | 109705 | 376229 | 126954 | 110504 |
| Sakai | 151151 | 143497 | 142760 | 114507 |
| TW14359 | 155589 | 134568 | 149038 | 127925 |
| 180 | 131237 | 144404 | 119660 | 128622 |
| Mean | 136920 | 199675 | 134603 | 120390 |

For all lineages, LCRs were biased toward the chromosomal terminus of replication (Ter) with the majority located between 2 Mbp – 3.5 Mbp (Fig. 1). The largest LCR identified was a 1. 4 Mbp inversion which was detected in Lineage Ic strain Z1615 (Fig. 1 and Supplementary Figure. S1). The average length of LCRs detected ranged between 109 – 376 Kbp depending on which reference strain was used for alignment with the largest LCRs detected within lineage Ic strains when aligned against lineage Ic reference strain 9000 (Table. 1 and Supplementary Figure. S3A). Mapping the chromosomal position of prophages within each reference genome further demonstrated that most LCRs were bounded by prophages (marked in red in comparison strain, Fig. 1). Furthermore, many of the LCRs identified had prophage Stx2c (ΦStx2c) as a boundary, particularly those occurring within Lineage Ic strains.

### LCRs map to repeated regions of homology on prophage

Mechanistically, chromosomal inversions typically involve recombination between inverted repeat regions of homologous sequences [22, 23]. As inversions were the dominant LCR identified in our analysis (Fig 1), we mapped the chromosomal position and direction of all homologous regions for each *E. coli* O157 strain (Fig. 2 and Table. S2). To avoid detection of the numerous IS elements present in *E. coli* O157 genomes [24] we restricted our analysis to regions that shared ≥ 98 % sequence homology, were ≥ 5000 bp and occurred in the chromosome with a frequency ≥ 2. Repeat regions were unequally distributed throughout the chromosome with a bias toward Ter and were conserved as inverted repeats at either side of Ter (Fig 2A). When each genome was subdivided into 1 Mbp domains, significantly more repeats were located within the 2 – 3 Mbp domain (p < 0.0001) adjacent to Ter than any other domain of the chromosome (Supplementary Figure. S3B). Significantly more repeats were also located within the 3 – 4 Mbp domain (p < 0.0001) adjacent to Ter than the 1 – 2 Mbp, 4 – 5 Mbp and 5 – 6 Mbp regions but not the 0 – 1 Mbp domain (p = 0.71). All repeat regions identified in the terminal half of the chromosome mapped within prophage (Fig 2B and Fig 2C) and specific combinations of these repeated regions matched the boundaries for identified LCRs. For example, specific recombination between regions 1a and 1b of Strain 9000 in Fig 2B would generate the LCR present in isogenic strain Z1767 and recombination between 2a and 2b would generate the LCR present in isogenic strain Z1615. These results indicate that homologous prophage sequences are hotspots for recombination resulting in the generation LCRs in *E. coli* O157 strains.

**Figure 2.**
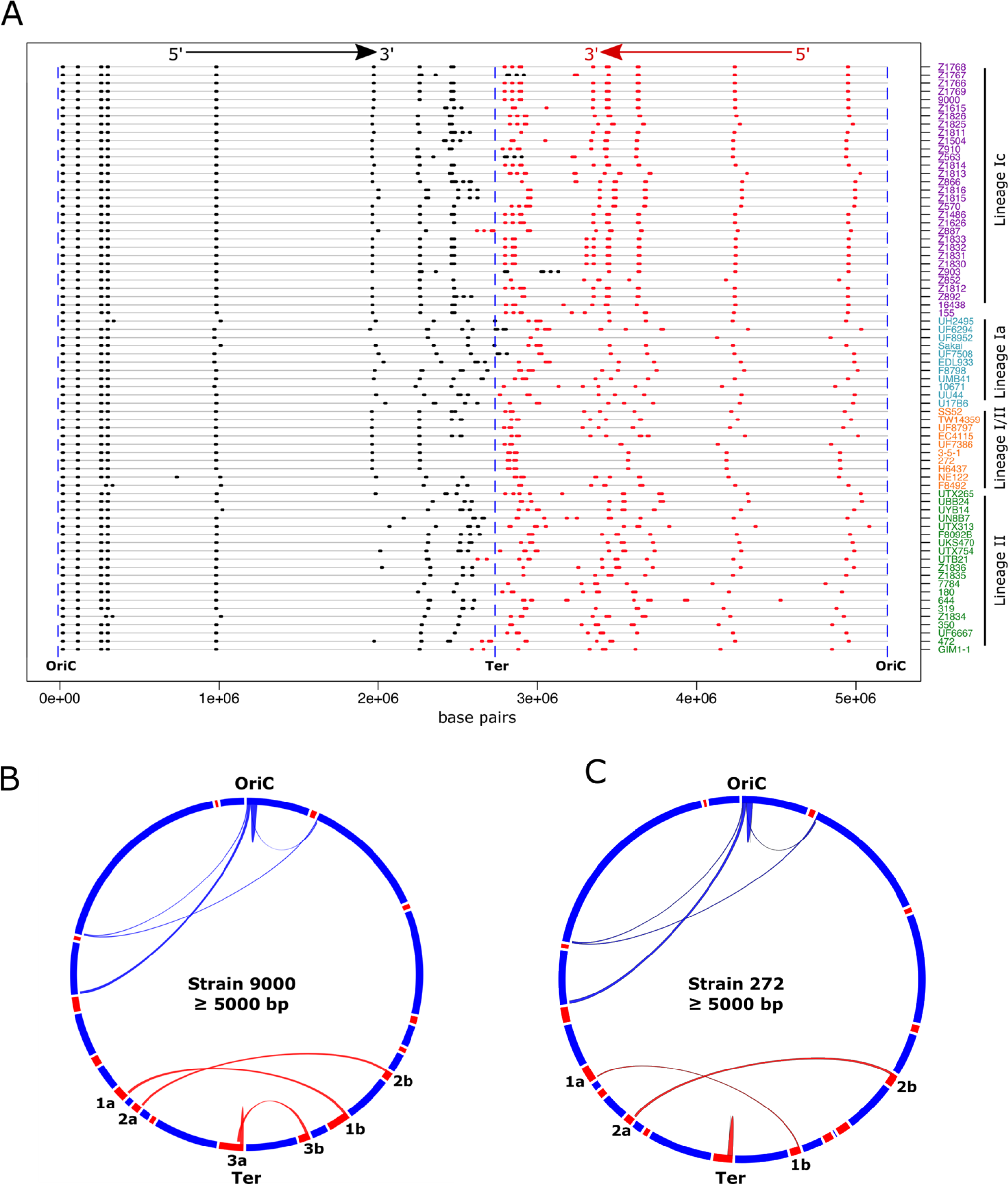
Mapping homologous regions (≥ 5000 bp) in *E. coli* O157. The loci of all regions of homology ≥ 5000 bp (black/red) that are present as ≥ 2 copies per genome are mapped on the chromosomes (grey line) of strains from Lineage 1c (purple), Lineage 1a (blue) Lineage I/II (orange) and Lineage II (green) (**A**). The directions 5’ – 3’ of homologous sequences relative to OriC are shown with black indicating the inverse direction to red. Circos plots for Lineage Ic strain 9000 (**B**) and Lineage I/II strain 272 (**C**) show paired regions of homology. Prophage loci (red blocks) are shown on the respective circular genome maps (blue). Paired homologous regions are joined by arches: Chromosomal (blue) and within prophage (red).

It was evident that specific combinations of inverted repeat regions were present in the different lineages and sub-lineages of *E. coli* O157 (Fig 2A). We reasoned that the frequency of recombinational events would be greater in strains with more homologous repeat regions and *vice versa*. Indeed, strains from Lineage Ic, in which the greatest number of LCRs were identified (Table. 1), had significantly more repeat regions > 5000 bp (p < 0.05) (Supplementary Figure S4A) and > 8000 bp (p < 0.0001) (Supplementary Figure S4B) than those from any other lineage. Conversely, Lineage I/II strains, in which only a single LCR was identified, had fewer homologous repeat regions ≥5000 bp than strains from any other lineage (Supplementary Figure S4A) and significantly less homologous repeat regions ≥ 8000 bp (p < 0.01) (Supplementary Figure S4B).

### LCRs underpin PFGE type expansion in Lineage Ic PT21/28 strains

In the United Kingdom, PT21/28 strains from Lineage Ic have arisen as the dominant PT associated with severe human infections over the last 20 years [11]. Based on standard pulsed-field gel electrophoresis (PFGE) typing methods, PT21/28 isolates have expanded from an initial 5 PFGE types (Profiles A - E, personal communication from Dr Lesley Allison Scottish *E. coli* reference laboratory-SERL) present in the UK in 1994 to >30 distinct PFGE profiles (Fig. 3A and Supplementary Figure. S5) by 2013 when PFGE was replaced by MLVA analysis. LCRs were shown to generate changes in the PFGE type of strain EDL933 [19], we therefore determined if LCRs also underpinned the PFGE type expansion seen in PT21/28 strains. We sequenced ten PT21/28 isolates by PacBio long-read sequencing that differed in PFGE type. Strains were selected from throughout the PT21/28 core SNP based phylogeny (Supplementary Figure. S5) and the dataset included two isolates with identical SNP addresses (Z910 and Z563; zero SNP differences in the core genome) but with distinct PFGE profiles.

**Figure 3.**
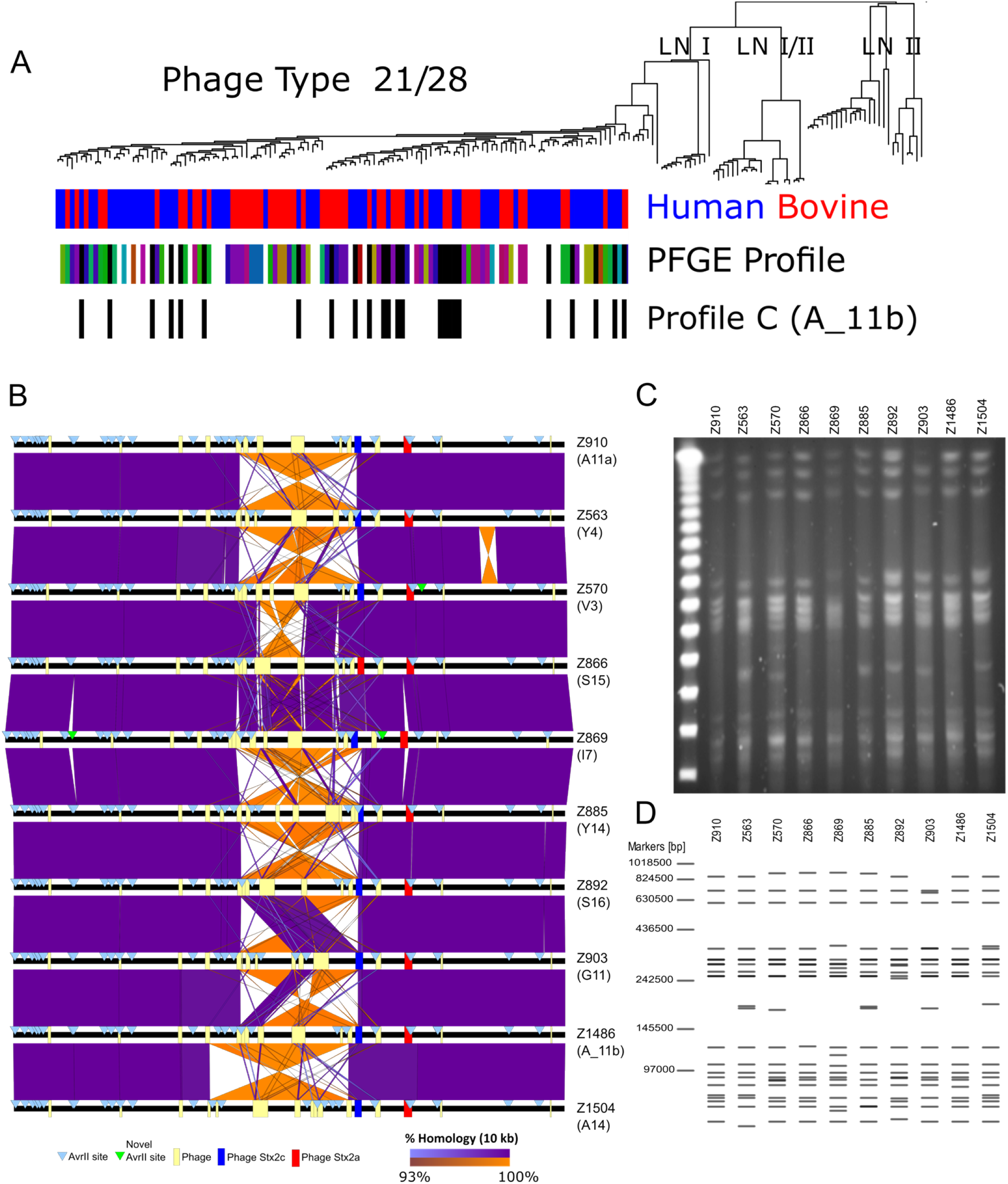
Distinct PFGE restriction patterns of *E coli* O157 PT21/28 strains are largely accounted for by LCRs. (**A**) Phylogenetic distribution of Lineage Ic PT21/28 strains. The source attribution, human (red) or bovine (blue) for each strain and PFGE variation across the lineage (coloured blocks) are shown. (**B**) Pairwise whole genome comparison of ten PT21/28 strains with different PFGE profiles. Whole genomes (black lines) are centred by the replication terminus (Ter) and loci of prophage (yellow boxes), Stx prophage (ΦStx2c;blue and ΦStx2a;red) and AvrII sites (blue triangles) are shown. Direct (purple) and inverted (orange) homology at a blast cut-off of 10,000 bp between strains are plotted. (**C**) PFGE profile of the ten selected PT21/28 strains following AvrII digestion. (**D**) *In silico* generated AvrII digestion pattern of the PacBio-generated sequences for each strain.

Sequence analysis showed that all ten strains differed by < 70 SNPs in their core genomes (Supplementary Figure. S5). Although examples of phage gain/loss (n = 2) were apparent, pairwise whole genome comparisons showed that LCRs were the dominant source of genomic variation at the macro scale (Fig. 3B). Reference laboratories specializing in STEC diagnostics in the UK used AvrII and/or XbaI restriction enzymes when determining the PFGE type for an isolate. When all AvrII restriction sites were mapped in each isolate (Fig. 3B) it was evident that the loci of most sites were strongly conserved. However significant strain variation in AvrII loci was observed within the Ter region of the chromosome that was associated with LCRs. For example, strains Z910 and Z563, which were identical at the core SNP level, differed by a single 1.2 Mbp chromosomal inversion that involved recombination with ΦStx2c and resulted in the repositioning of four AvrII sites. Additional sequences containing AvrII sites present in some strains but not in others were identified (Fig. 3B) however these were rare. The majority of AvrII loci variation and therefore PFGE type variation was generated by LCRs.

To confirm that the variation in AvrII loci generated by LCRs observed in our PacBio assemblies matched the actual chromosome configuration of each isolate we determined the PFGE profile for each strain after AvrII restriction digestion (Fig. 3C) and compared it to *in silico* AvrII digests of their respective Pac-Bio assemblies (Fig. 3D). Both *in vivo* and *in silico* AvrII digestion patterns were matched for 9/10 strains analysed confirming the presence of those LCRs identified by PacBio long-read sequencing and the rearrangement of AvrII loci by these LCRs to generate different PFGE types. The exception was strain Z892 in which an unexplained digestion product was present after *in vivo* digestion that was not predicted from the PacBio sequence.

Based on these results we propose that the majority of the PFGE variation amongst PT21/28 strains, as depicted in Fig. 3A and Supplementary Figure. S5, is generated by LCRs. It was also evident from the PFGE analyses that the strains cultured under these laboratory conditions had the majority of their genomes in a single confirmation as there was no evidence of weak secondary bands in the gel restriction patterns (Fig. 3C). Of note, the most frequently occurring PT21/28 strain PFGE profile was type ‘C’ later defined as profile A_11b (Fig. 3A and Fig. S5). Phylogenetically, this specific profile re-occurs throughout the sub-lineage indicating that it is likely an ancestral confirmation or strains can repeatedly return to this chromosome conformation.

### *In vivo* occurrence of LCRs during host colonisation

Previously, we carried out a series of published and unpublished *in vivo* cattle colonization studies focused on *E. coli* O157 strain 9000 [25, 26]. To determine if LCRs are present during animal colonization we compared isolates collected from two separate colonization studies by PacBio long-read sequencing and AvrII PFGE profiling. This isolate set were all derivatives of the original wildtype strain 9000 and included inoculum and recovered isolates (Supplementary Table S1).

Pairwise whole genome comparisons of strains 9000 and Z1615 from Trial 1 (Fig. 4A) showed a 1.4 Mbp inversion had occurred in derivative strain Z1615 relative to strain 9000. As outlined in Fig. 2B the boundaries of this LCR mapped to large inverted repeat sequences within prophage located either side of Ter. Distinct PFGE profiles were observed for strains 9000 and Z1615 following AvrII digestion, each matched their respective *in silico* AvrII digestion profiles (Supplementary Figure S6A) and confirmed the presence of the LCR identified in Z1615. No evidence of secondary bands diagnostic of Z1615 chromosomal conformation in the PFGE profile of strain 9000 were observed indicating this LCR occurred or was selected during colonization to generate strain Z1615. To determine how frequently this LCR occurs we analysed a further eleven recovered isolates from two experimental trials (Trial 1 and Trial 3) in which strain 9000 was the inoculum by PFGE (Supplementary Figure S6B). Isolates were collected from a number of different animals and dates (Supplementary Table S1). Three additional isolates of the 11 tested matched the PFGE profile of Z1615 indicating an *in vivo* selection for this LCR in the bovine host.

**Figure 4.**
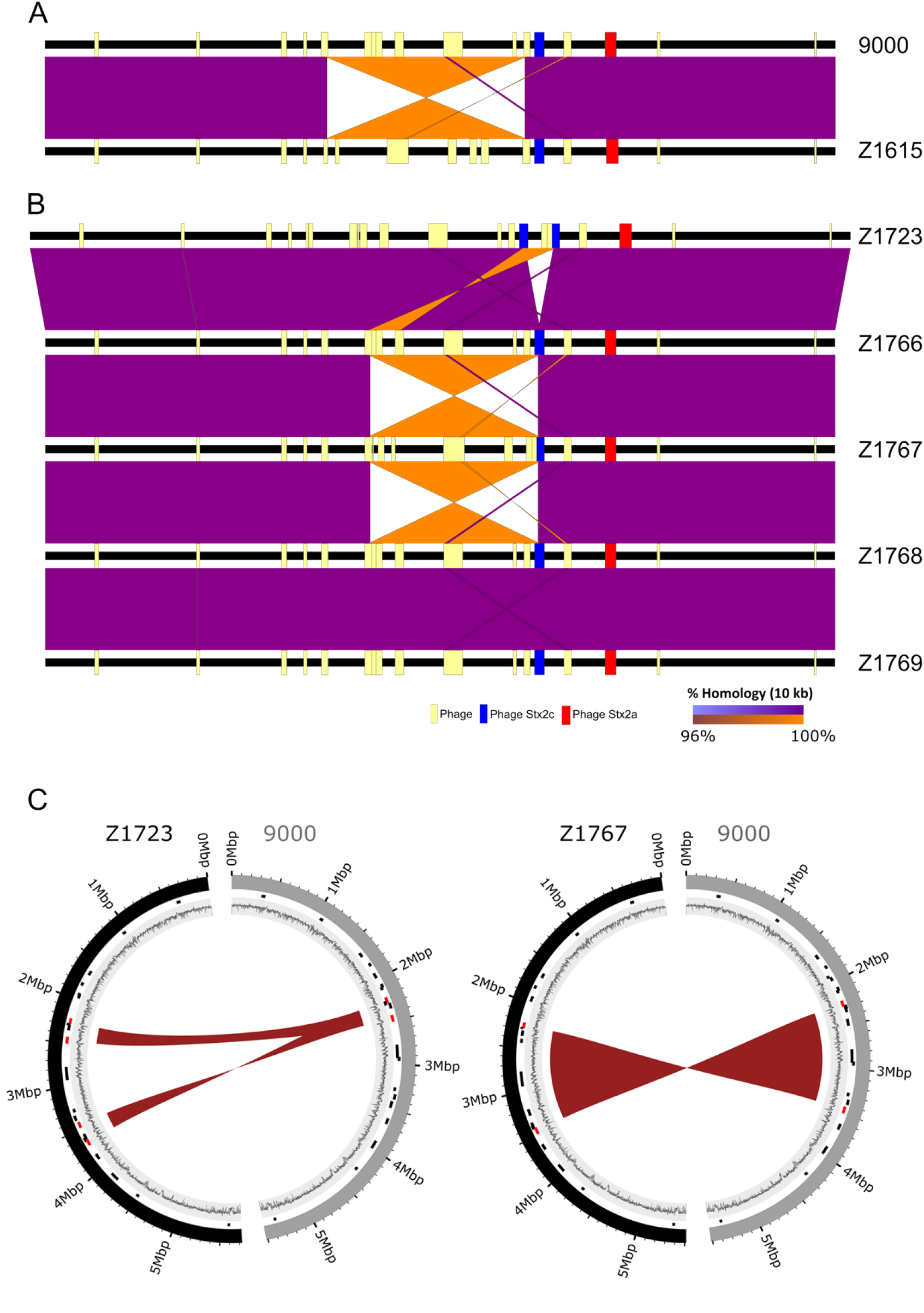
Detection of LCRs in *E. coli* O157 PT21/28 strain 9000 variants analysed from cattle colonization studies. Pairwise whole genome comparisons of strains from Trial 1 (**A**) and Trial 2 (**B**) are shown with direct (purple) and inverted (orange) homology at a blast cut-off of 10,000 bp between strains. Whole genomes (black lines) are centred by the replication terminus (Ter) and the loci of prophage (yellow boxes) and Stx prophage (ΦStx2c;blue and ΦStx2a;red) in each strain are mapped. (**C**) Circos plots showing the identified 220 kbp duplication in Z1723 (left) and 1.2 Mbp inversion in Z1767 (right) relative to progenitor strain 9000. Outer ring: Strain 9000 (grey) and LCR derivatives Z1723 and Z1767 (black); Middle ring: Loci of prophage (black) and prophage a LCR boundaries (red); Inner ring: GC content.

Two additional LCRs were identified from the five isolates examined from Trial 2 (Fig. 4B) both of which involved recombination with the Stx2c prophage (ΦStx2c). A 220 kbp inverted duplication was identified in strain Z1723. The duplicated region was flanked by repeat sequences from within prophage located at 2.2 Mbp and 2.4 Mbp (Supplementary Table S2) relative to OriC and inserted into ΦStx2c (3.4 Mbp) bisecting the Stx2c prophage (Fig. 4B and Fig. 4C). A second 1.2 Mbp inversion was identified in strain Z1767 that also involved recombination between repeat sequences within the same prophage located at 2.2 Mbp and ΦStx2c (Fig. 4B and Fig. 4C). PFGE analysis confirmed the presence of the LCRs in Z1723 and Z1767 (Supplementary Figure. S6A).

### Real-time occurrence of LCRs during *in vitro* laboratory culture

We investigated if LCRs could be generated and detected in real-time following standard laboratory culture of bacteria in LB media. To increase the sensitivity of detection we applied both Oxford Nanopore Technologies (ONT) long-read sequencing and optical mapping to detect LCRs in strains from animal colonization Trials 1 and 2.

The wildtype parental strain 9000 was first sequenced using ONT and searched for reads that aligned to the LCRs identified in variant strains Z1615, Z1767 or Z1723. Aligning strain 9000 reads to the Z1615 genome, a total of 5 reads were found that matched the identified 1.4 Mb inversion boundary at 1.95 Mb relative to OriC and a single read that matched the inversion boundary at 3.35 Mb. These reads were abundant at approximately 2 % and 0.33 %, respectively, of the total reads across the same region that mapped directly to strain 9000. Similarly aligning 9000 reads to the Z1767 genome, a single read (0.4 % abundance) was found that matched the 1.2 Mb inversion boundary at 2.25 Mb relative to OriC and three reads (1.2 % abundance) that matched the inversion boundary within ΦStx2c at 3.45 Mb. No reads were found that mapped to the 220 kb duplication in Z1723.

Next we analysed strains Z1723 and Z1767 using Bionano Irys optical mapping (Figure 5 and Supplementary Figure S6) to identify additional LCRs that occur during growth in LB medium. Cultures of each strain were started from single colonies and chromosomes were extracted during late exponential phase cultures (OD600 = 0.7). Structural variant (SV) analysis was performed to detect all novel genome restriction maps within the cultured populations of Z1723 and Z1767 that did not map directly to an *in silico* generated map of the parental strain 9000 reference genome (Fig. 5).

**Figure 5.**
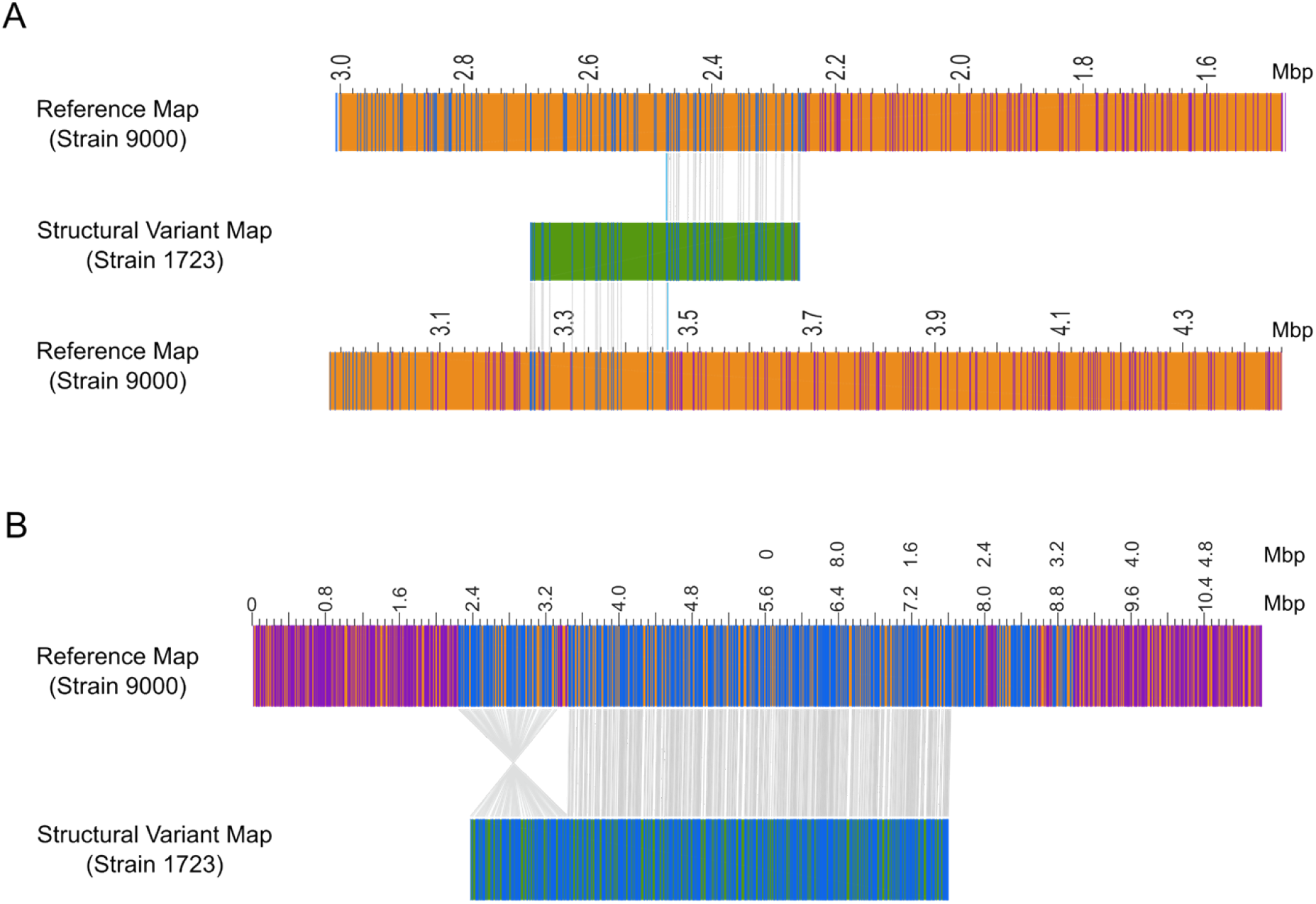
Optical mapping of *E. coli* O157 PT21/28 strain 9000 variant Z1723. Structural variant (SV) analysis identified a 220 Kb duplication (**A**) and 1.2 Mb inversion (**B**) in the population of Z1723 relative to the reference strain 9000. The genome map (orange) of reference strain 9000 and each Z1723 structural variant (green) are shown. Paired restriction sites (blue lines) are aligned between the reference and variant maps (grey lines). Unpaired restriction sites (purple lines) outside aligned regions are also shown. The SV map containing the 220 Kb duplication has been aligned to two reference strain 9000 genome maps to demonstrate the hybrid composition of the map containing ΦStx2c at 3.4 Mb and an inverted 220 Kb duplicated region originating from between 2.2 and 2.4 Mb.

Optical mapping showed that both strains had mixed population structures when cultured *in vitro*. SV analysis confirmed the same 220 kb inverted duplication was present in the Z1723 population that was identified by PacBio sequencing and PFGE (Fig. 5A). This hybrid structural variant mapped 5’ – 3’ between 2.24 – 2.46 Mb and 3’ – 5’ between 3.26 – 3.46 Mb to Strain 9000 further confirming the presence of the inverted duplication within ΦStx2c at 3.4 Mb. A 1.2 Mbp inversion relative to strain 9000 (Fig. 5B) was also identified in Z1723. This inversion matched the 1.2 Mbp inversion seen in strain Z1767 (Fig. 4B) with boundaries in prophage located at 2.2 Mbp and 3.4 Mbp (ΦStx2c). PFGE analysis of two separate Z1723 freezer stocks (Supplementary Figure S6A) shows that the 220 kbp inverted duplication is the dominant genome conformation present with no evidence of secondary bands indicative of the Z1767 inversion. We therefore assume that the 1.2 Mbp inversion detected in the Z1723 population by optical mapping is a minority population below the limit of detection by PFGE.

SV analysis of Z1767 identified the expected 1.2 Mbp inversion relative to strain 9000 (Supplementary Figure S7A) as determined from Pac-Bio sequencing and identified a novel 140.5 kbp inverted duplication within the cultured population (Supplementary Figure S7B). The duplicated region spanned 2.1 – 2.24 Mbp relative to OriC and was flanked by prophage sequence (2.2 Mbp) and an IS66 sequence located within the O-Island 48 [27]. This duplicated region also inserted in an inverted orientation within the Stx2c prophage further highlighting ΦStx2c as a hotspot for recombinational events leading to LCRs.

### Changes in bacterial gene expression and phenotypes associated with LCRs

Using the structural variants of strain 9000 (Z1723, Z1767, Z1615) generated during *in vivo* colonization we examined if the identified LCRs impacted strain phenotypes. The global transcriptomes of strain 9000 and each structural variant strain (Z1723, Z1767, Z1615) were first compared by RNAseq for two growth conditions: nutrient rich LB medium and minimal M9 medium. PCA analysis showed there was little discernible difference between the transcriptomes of each strain when cultured in LB (Supplementary Figure S8A) however the transcriptome of strain Z1723, containing a 220kbp inverted duplication, was distinct from strain 9000 and the other variants in M9 (Supplementary Figure S8B). Differential changes in gene expression were modest (Supplementary Table S3) although a gene dosage effect was apparent across the region of duplication with an increase in expression observed for 66 of the duplicated genes when mapped to the genome of WT strain 9000 (Fig. 6A). There was also a marked effect on the expression of genes within the Stx2a prophage (ΦStx2a) rather than the Stx2c prophage (ΦStx2c) into which the 220 kbp duplication had inserted (Fig. 6A).

**Figure 6.**
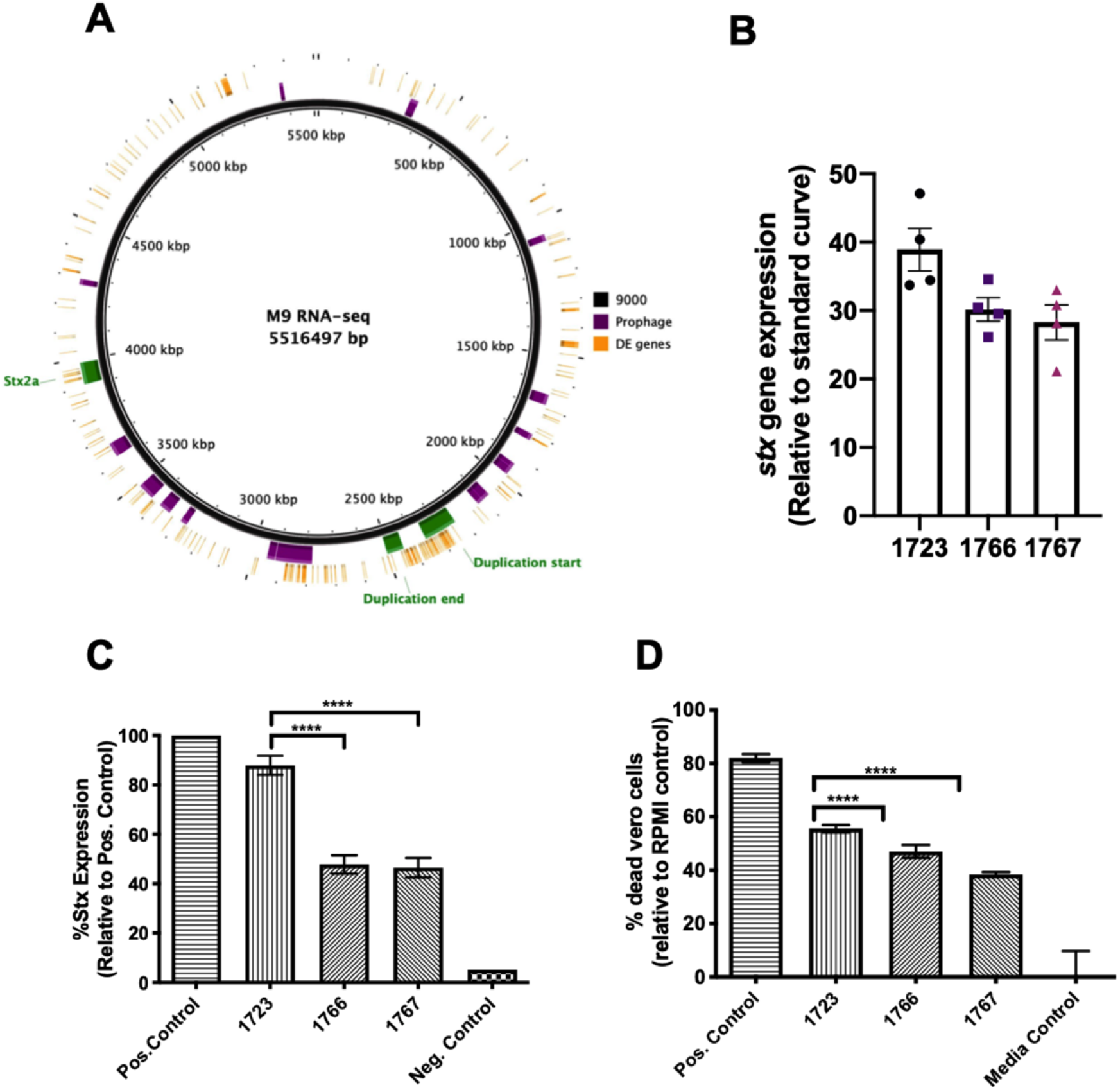
Shiga toxin expression, production and toxicity of Strain 9000 structural variants. The chromosomal location of all differentially expressed genes in Z1723 (orange bars) are mapped to the reference strain 9000 genome (**A**). Prophage (red blocks), the 220 kb duplication and ΦStx2a regions are highlighted. Expression of stx2a (**B**) total Stx toxin production (**C**) and Vero cell toxicity Stx (**D**) was measured for Trial 2 strains Z1723, Z1766 and Z1767 in M9 media. Mean values +/− SEM of four biological replicates (n = 4) are shown for each assay. * p ≤ 0.05; ** p ≤ 0.01; *** p ≤ 0.001; ****p ≤ 0.0001

As Stx2 toxin is the primary virulence factor of *E. coli* O157 strains leading to HUS, we tested if the observed differential transcription within ΦStx2a in Z1723 affected Stx2a expression, production and activity compared with other structural variants. For each phenotype Z1723 was compared with Trial2 variants Z1766 and Z1767. Strains 9000 and Z1615 were excluded due to the previously documented [25] inactivation of the *stx2a* gene by an IS element, IS629. Expression of *stx2a* was increased in Z1723 compared to both Z1766 and Z1767 (Fig. 6B) and this manifested as a significant increase in total Stx2 toxin (Fig. 6C) and cytotoxic killing of Stx2 susceptible Vero cells (Fig. 6D).

We have previously shown that lysogeny with Stx2 prophages negatively regulates the LEE type III secretion system (T3S) [28] and demonstrated that a large duplication may have influenced the fitness of two closely related outbreak strains [29]. As the 220 kbp duplication in Z1723 interrupted ΦStx2c and increased expression of ΦStx2a genes we examined T3S and assessed the competitive fitness for strains Z1723, Z1766 and Z1767 (Fig. 7). Transcriptional *gfp* fusions to the LEE master regulator, *ler*, and LEE4 encoded *sepL* were introduced into each strain and expression was monitored in MEM-HEPES medium (OD600 = 0.8). Expression of both *ler* and *sepL* was decreased in Z1723 compared to Z1766 and Z1767 (Fig. 7A). There was also marked difference in the levels of the T3S secreted protein, EspD, which could not be detected in the culture supernatant of Z1723 (Fig. 7B).

**Figure 7.**
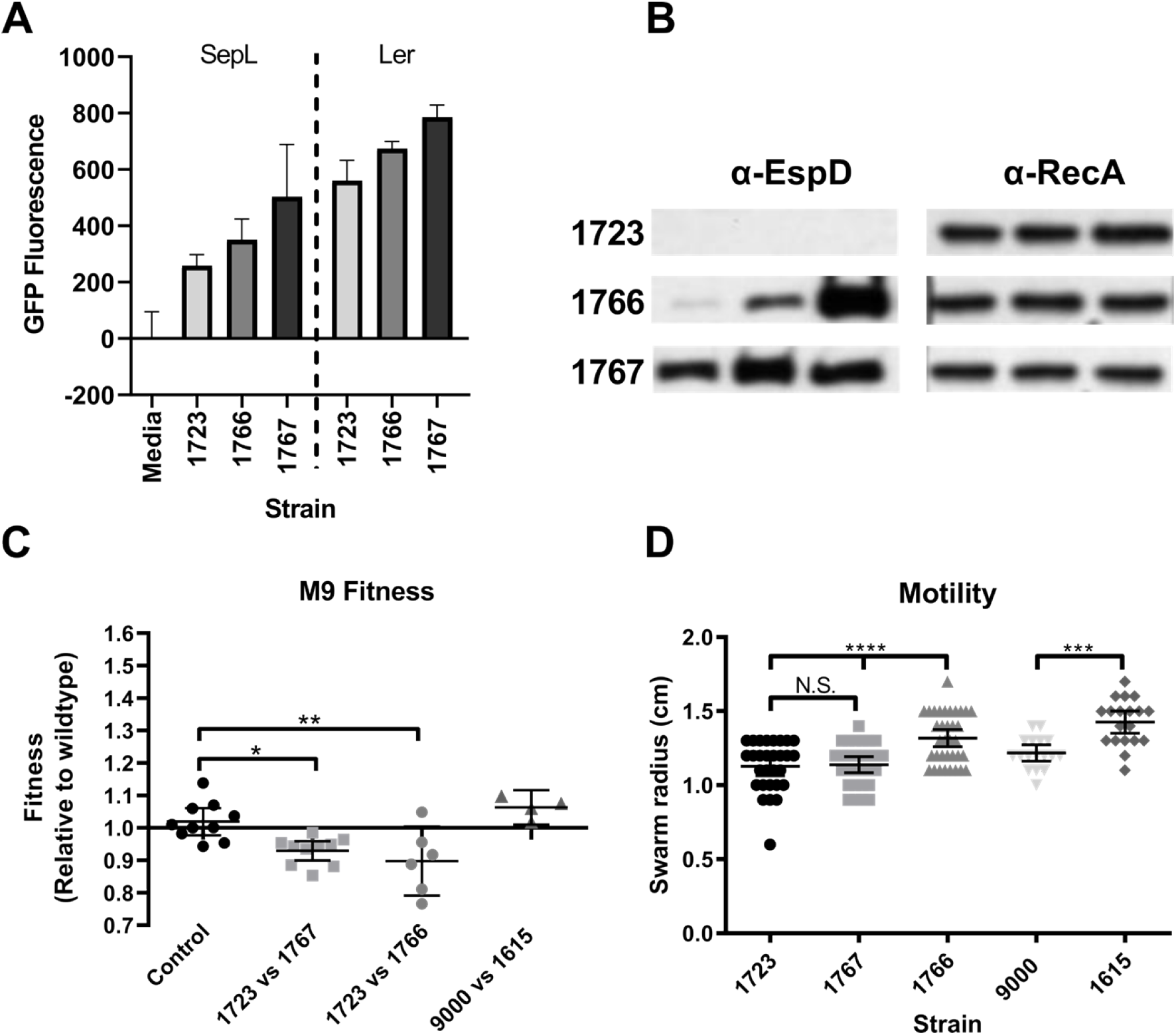
Type III secretion, competitive fitness and motility phenotypes of strain 9000 structural variants. **(A)** Expression of the LEE master regulator *ler* and LEE4 chaperone *sepL* was measured by Gfp reporter fusions (n = 3). **(B)** Detection of the LEE effector EspD in the culture supernatants of each strain by Western blot (n = 3). Corresponding cellular RecA levels were used as a control. (**C**) Competitive fitness of strains after 24 h co-culturing in M9 media (n = 6). **(D)** motility of strains after 6 h on Tryptone swarm plates (n = 20). Mean values +/− SEM are shown for each assay. * p ≤ 0.05; ** p ≤ 0.01; *** p ≤ 0.001; ****p ≤ 0.0001

The competitive fitness of strains from Trial 1 (9000 and Z1615) and Trial 2 (Z1723, 1766, 1767) was assessed by paired co-culturing in M9 media. In M9 media Z1723 significantly outcompeted the structural variants Z1766 and Z1767 as mean fitness indices (f.i.) of 0.89 and 0.93 were recorded, respectively (Fig. 7C) compared with control, f.i. = 1. No significant difference in fitness was observed between trial 1 strains in M9 (Fig. 7C). Finally, we measured the motility of strains 9000 and each structural variant on tryptone swarm plates (Fig. 7D). For strains isolated from calf trial 2 no difference in motility between Z1723 and Z1767 was observed however Z1766 was significantly more motile than both variants. Z1615 from calf trial 1 was also significantly more motile than WT strain 9000. These data provide evidence that LCRs can impact important *E. coli* O157 phenotypes involved in host colonisation and disease.

## Discussion

Phenotypic heterogeneity within isogenic populations of microorganisms is used as a ‘bet-hedging’ survival strategy to cope with sudden fluctuations in environmental conditions and can lead to a division of labour between individuals that raises group fitness [30–33]. We have demonstrated that *E. coli* O157 can generate such heterogeneity through LCRs occurring between homologous prophage sequence *in vivo* and *in vitro*. As originally demonstrated for *E. coli* O157 strain EDL933 [19] the LCRs we have now documented across the serogroup are bounded by specific prophages clustered towards the terminus of the genome. Chromosomal inversions involving the Ter region that lead to replichore imbalance can stall or stop replication forks and induce SOS [34]. Due to the spatial distribution of prophages involved in LCRs, the main large inversions we have identified do not generate major changes in replichore size. However even minor changes could impact growth rate and phenotypes as seen with the LCR specific phenotypes identified in this study affecting virulence gene expression, fitness and motility.

Large prophage homologous repeats (> 5000 bp) were identified at the boundaries of LCRs which provide ample sequence substrate for recombination. In addition to RecABCD-mediated recombination, *E. coli* O157 strains also carry multiple λ-like phage, including Stx phage, many of which encode their own Rad52-like recombinase enzymes such as Redβ [35–37]. Whether the formation of LCRs in *E. coli* O157 strains is host or phage mediated is unknown. Irrespective of the recombination system involved, the generation of LCRs would require a double-strand break (DSB) in one or more of the phage at their boundaries. It is interesting to speculate that double-strand breaks (DSBs) within phage are a primary driver of recombinational repair in bacteria via the SOS response that is also required for prophage-based expression of Stx. Strains of *E. coli* O157 PT21/28 constitutively express Stx2 [25] and therefore the rate of occurrence of DSBs, RecA-mediated Stx expression and LCR formation may be interconnected.

The Stx2c-encoding prophage was shown to be present across the different *E. coli* O157 lineages without much variation compared to Stx2a encoding prophages [9, 11]. In the present study it is a primary architect of many of the LCRs and as such may be subject to positive selection. One structural variant of PT21/28 strain 9000 was a duplication from one side of the chromosome inserted into the Stx2c terminase gene region on the other side of the genome. This was of particular interest as this large region of duplicated homology would stimulate inversions and also recombination resolving back to the original confirmation. A similar duplication has been sequenced in two closely related strains associated with sequential *E. coli* O157 outbreaks at a single restaurant [19]

The ONT long-read sequencing and optical mapping results provide evidence that LCRs are continuously generated at very low levels. The estimation from the ONT long-read sequencing of strain 9000 was between 1 – 2 % of the population when cultured in LB. For these specific LCRs to be detectable during animal colonization indicates that they have been selected under the *in vivo* conditions of the animals intestinal tract. For example, in colonisation experiments, the input strain 9000 confirmation (profile C/A_11b) was recovered in 8/12 isolates, with the remaining four having the large 1.4 Mb inversion as determined by PFGE (Fig. S6B). Intriguingly, the highest excretion level in that experiment was associated with an animal from which a strain with the inverted confirmation was recovered. Currently, there is no simple way to quantify the proportions of the confirmations under specific conditions, with the exception of optical mapping for the isolates cultured in the laboratory.

As further support for these processes in cattle, extensive surveys of *E. coli* O157 in cattle herds [38, 39] determined that while the majority of isolates in any specific herd exhibit the same PFGE pattern, there are isolates with different profiles yet the same phage type (PT) [40, 41]. A recent study of persistent Lineage I strains isolated on a single farm also demonstrated that a 47.7 kbp deletion was a significant genomic difference between two of the strains [42]. We show that LCRs are the likely cause of the observed PFGE profile type expansion amongst PT21/28 bovine isolates in the UK. There has been one previous report of multiple deletions occurring during *E. coli* O157 colonisation of cattle, generating multiple PFGE types [43].

LCRs have been observed in a number of bacterial genera, including *Campylobacter*, *Yersinia*, *Staphylococcus* and *Salmonella* [22, 44–47]. For inversions the gene content and copy number is maintained but the prophage boundaries do change in composition and this could have an impact on prophage gene expression or the regulatory networks that they are part of [1]. A clear example of this was shown for *Campylobacter* where in one orientation the inversion completes an active prophage and in turn that provides resistance to certain infecting phages [44]. For *E. coli* O157 strain 9000 structural variants we measured a number of expression and phenotype changes, including motility and growth rate for variants with inversions. The most obvious differentials were present in the variant with a 220 kbp duplication. This included an increase in Stx expression, production and toxicity and a reduction in type 3 secretion. Our previous research has shown that Stx2a prophage integration into different *E. coli* backgrounds led to a repression of T3S, potentially via the CII protein [28]. Such cross-regulation would offer one pathway resulting in the concomitant reduction in T3S in the strain with the duplication.

## Conclusions

We describe the first systematic genome structure comparison of strains across the main lineages *Escherichia coli* O157. LCRs, predominantly large inversions, were a common source genomic variation and appear to be generated by recombination between homologous prophage sequences. Importantly, we show that LCRs are generated during animal colonisation and laboratory culture and demonstrate that specific LCRs are associated with phenotypic changes. By definition, phenotypic heterogeneity is the occurrence of individuals within a genetically identical population that stochastically develop phenotypes of varying fitness within a homogenous environment [30, 32]. With the work presented here and that in other genera, it is evident that genome structural variants are a way to generate phenotypic heterogeneity in a clonal bacterial population and that relevant sub-populations can then be selected as conditions change in particular environments making it an important population survival strategy.

## Materials and Methods

### Bacterial strains and culture conditions

Bacterial strains and plasmids used in this study are listed in Table. S1. Bacteria were cultured in Luria-Bertani (LB) broth or M9 minimal media (Sigma-Aldrich) supplemented with 0.2% glucose, 2 mM MgSO_4_ and 0.1 mM CaCl_2_. For TTSS expression bacteria were cultured overnight in LB and then inoculated into minimal essential medium (MEM)-HEPES (Sigma-Aldrich) supplemented with 0.1% glucose and 250 nM Fe(NO_3_)_3_. Antibiotics were used at the following concentrations when required: Chloramphenicol (50 μg/ml), Mitomycin C (2 μg/ml), Nalidixic acid (50 μg/ml).

### PacBio Long-read sequencing

A total of 72 whole genome sequences, generated by PacBio long-read sequencing, were used for analysis in this study. The sequences of 31 strains were determined for this study and the remaining 41 were publicly available in the National Centre for Biotechnology Information (NCBI) database (Table S1).

Sequencing of the 31 isolates was conducted using a PacBio RS II long-read sequencing platform and carried out at the U. S. Department of Agriculture sequencing core facility in in Clay Center, Nebraska, USA. Qiagen Genomic-tip 100/G columns and a modified protocol, as previously described [48], were used to extract high molecular weight DNA. Using a g-TUBE (Corvaris), 10 μg of DNA was sheared to a targeted size of 20 kb and concentrated using 0.45x volume of AMPure PB magnetic beads (Pacific Biosciences). Following the manufacturer’s protocol, 5 μg sheared DNA and the PacBio DNA SMRTbell Template Prep kit 1.0 were used to create the sequencing libraries. A BluePippin instrument (Sage Science) with the SMRTbell 15–20 kb setting was used to size select 10 kb or larger fragments. The library was bound with polymerase P5 and sequencing was conducted with the C3 chemistry and the 120 min data collection protocol. Individual libraries were constructed from some of the strain DNA preparations described above using an Illumina Nextera XT DNA sample preparation kits with appropriate indices tags according to the manufacturer’s instructions (Illumina Inc., San Diego, CA). The libraries were pooled together and run on an Illumina MiSeq DNA sequencer (Illumina Inc., San Diego, CA). The genome of each strain was sequenced to a targeted depth of 50X coverage.

### Genome assembly and annotation

SMRT analysis was used to generate a FASTQ file from the PacBio reads, which were then error-corrected using PBcR with self-correction [49]. The Celera Assembler was used to assemble the longest 20× coverage of the corrected reads. The resulting contigs were improved using Quiver [50] and annotation was conducted using a local instance of Do-It-Yourself Annotator (DIYA) [51]. Geneious (Biomatters) was used to remove duplicated sequence from the 5′ and 3′ ends to generate the circularized chromosome. To correct PacBio sequencing errors (homopolymers and SNPs), Illumina reads were mapped to the Quiver polished chromosome using Pilon [52]. Then, both PacBio and Illumina reads were mapped to the Pilon-generated chromosome using Geneious Mapper. Additional sequencing errors were identified and corrected by manual editing in Geneious, resulting in a finished closed circularized chromosome. OriFinder was used to determine the origin of replication [53] and the chromosome was reoriented using the origin as base number one. Prophage regions were identified as described previously [9] using PHASTER [54].

### MinION sequencing and SV read detection

Strain 9000 was sequenced by Oxford nanopore technologies MinION sequencing. High molecular weight genomic DNA was extracted from strain 9000 grown in LB (OD600 = 0.7) by standard phenol:chloroform extraction [55]. Genomic DNA was purified using Qiagen G100 Genomic Tips (Qiagen) with minor alterations including no vigorous mixing steps and final elution in 100μl of nuclease free water and quantified using a Qubit and the HS (high sensitivity) dsDNA assay kit (Thermofisher Scientific), following the manufacturer’s instructions. Library preparation was performed using the Ligation kit SQK-LSK109 (Oxford Nanopore Technologies). The prepared libraries were loaded onto a FLO-MIN106 R9.4.1D flow cell (Oxford Nanopore Technologies) and sequenced using the MinION (Oxford Nanopore Technologies) for 72 h. Data produced in a raw FAST5 format was basecalled and de-multiplexed using Guppy v3.2.4 using the FAST protocol (Oxford Nanopore Technologies) into FASTQ format.

To identify if the Nanopore sequenced strain 9000 contained reads supporting multiple isoforms of the chromosome. Minimap2 v2.17 [56] and Samtools v1.7 [57] was used to align the Nanopore reads (removing secondary aligning reads) to samples Z1615, Z1723 and Z1767 each representing a different chromosomal isoform. Using Samtools v1.7 [57] and Bedtools v2.29.2 [58] reads were identified at either end of the each of the 5’ and 3’ breakpoints identified in those conformations. The number of reads that crossed each end of the 5’ and 3’ breakpoints for both conformations was calculated again using Samtools v1.7 [57] and Bedtools v2.29.2 [58].

### Whole genome comparisons

Pairwise whole genome alignments were conducted with Easyfig [59] as described previously [9]. Genome .gbk files were modified so that prophage were represented as coloured blocks. AvrII restriction sites were identified in selected genomes using UGENE [60] and their loci were added to the respective genome .gbk files. Pairwise whole genome alignments between reference genomes from each lineage (9000, Sakai, TW14359 and 180) and each genome were performed using blastn [61] with the following parameters (-evalue 1e-10 -best_hit_score_edge 0.05 -best_hit_overhang0.25 -perc_identity 70 -max_target_seqs 1 -outfmt 6). From the resulting alignment files, LCRs were identified within each genome by filtering all inverted homologous regions ≥ 50,000 bp relative to each reference strain.

### Mapping homologous regions

Homologous regions within each genome were identified using blastn [61]. Blastn was performed on each individual genome using the same genome sequence as both reference and query with the following parameters (-evalue 1e-10 -best_hit_score_edge 0.05 -best_hit_overhang 0.25 - perc_identity 98 -max_target_seqs 1 -outfmt 6). Homologous regions that satisfied three conditions simultaneously were extracted from the blast output: (1) Homologous regions were ≥ 5000 bp (2) homologous regions ≥ 5000 bp were present in the genome at a frequency ≥ 2 (3) homologous regions were located before and after *dif* (terminus of replication). Equivalent analysis was repeated to determine homologous regions ≥ 8000 bp. Significant differences in the total number of repeats detected between lineages and the bias of repeat regions toward Ter was determined by one-way ANOVA with Dunnetts multiple comparisons test.

Circos plots [62] were used to visualise linked regions of homologous sequence within the genomes of selected strains. Custom circos input files were generated in which the data matrix was modified such that each circular genome was divided at prophage boundaries. Linked homologous regions and their sizes were determined using BLAST scores derived when querying a selected genome sequence to itself. Only BLAST hits with ≥ 98 % sequence homology and that were ≥ 5000 bp in length were included in the data matrix of circos input files. Within Circos plots the width of linked segments is proportional to the length of BLAST hits. Circos does not exactly map homology hits to linked chromosomal/prophage regions, instead connecting segments originate and end at the earliest available location within the linked region.

### Phylogeny of 72 strains and PT21/28 strains

A core gene alignment was extracted from the fully assembled and annotated PacBio genomes of all 72 strains using ROARY [63] with parameters (-e -n -r -s -ap). The extracted multiple alignment was used to Maximum-liklihood phylogenetic trees FastTree [64] (-gtr) and trees were visualised with iTOL [65]. To determine the phylogenetic relationship of PT21/28 strains high quality illumine sequencing reads were mapped to the reference STEC O157 strain, Sakai (GenBank accession BA000007), using Burrows-Wheeler Aligner – Maximum Exact Matching (BWA MEM (v0.7.2)) [66]. The sequence alignment map output from BWA were sorted and indexed to produce a binary alignment map (BAM) using Samtools (v1.1) [67]. Genome Analysis Toolkit (GATK v2.6.5) was then used to create a variant call format (VCF) file from each of the sorted BAMs, which were further parsed to extract only SNP positions of high quality (mapping quality (MQ) > 30, depth (DP) > 10, variant ratio> 0.9). Hierarchical single linkage clustering was performed on the pairwise SNP difference between all isolates at descending distance thresholds (Δ250, Δ100, Δ50, Δ25, Δ10, Δ5, Δ0) [68]. SNP alignments were created tolerating positions where >80% of isolates had a base call with regions of recombination masked using Gubbins v2.0.0 [69]. Maximum likelihood phylogenies were computed using IQ-TREE v2.0.4 [70] with the best-fit model automatically selected and near zero branches collapsed into polytomies.

### <u>P</u>ulsed-<u>f</u>ield <u>G</u>el <u>E</u>lectrophoresis

All strains analysed by PFGE were cultured in LB or M9 medium at 37°C overnight with agitation. Genomic DNA was purified using the CHEF Bacterial Genomic DNA Plug Kit (Bio-Rad) according to manufacturer guidelines. DNA restriction digestion with AvrII (BlnI) (Takara) and subsequent PFGE was done according to the PulseNet O157 guidelines [71], using a CHEF-DR III system. *In silico* AvrII (BlnI) restriction digests of selected genomes was carried out in CLC Genomics Workbench (Qiagen).

### RNA sequencing

Total RNA was extracted from three biological replicates of strains 9000, Z1615 Z1723, Z1767 using mirVana™ miRNA Isolation Kit (ThermoFisher) according to manufacturer guidelines.

Strains were cultured in either LB or M9 media to OD_600_ = 0.7. Ribosome depletion, cDNA library preparation and Illumina sequencing was carried out by Vertis Biotechnologie AG (Freising, Germany). Total RNA samples were purified and concentrated using the Agencourt RNAClean XP kit (Beckman Coulter Genomics) and the RNA integrity was assessed by capillary electrophoresis. Ribosomal RNA molecules were depleted using the Ribo-Zero rRNA Removal Kit for bacteria (Illumina). The ribodepleted RNA samples were first fragmented using ultrasound (4 pulses of 30 s each at 4°C) and oligonucleotide adapters were then ligated to the 3’ end of the RNA molecules. First-strand cDNA synthesis was performed using M-MLV reverse transcriptase and the 3’ adapter as primer. The first-strand cDNA was purified and the 5’ Illumina TruSeq sequencing adapter was ligated to the 3’ end of the antisense cDNA. The resulting cDNA was PCR-amplified to about 10-20 ng/μl using a high-fidelity DNA polymerase. The cDNA was purified using the Agencourt AMPure XP kit (Beckman Coulter Genomics) and was analyzed by capillary electrophoresis. Purified cDNA was pooled and sequenced on an Illumina NextSeq 500 system using 75 bp read length. RNA-sequencing reads were mapped to the strain 9000 reference genome (CP018252.1) using STAR 2.7.0e [72] with the following parameters (--quantMode GeneCounts and --sjdbGTFfeatureExon CDS). Prior to read mapping the reference strain 9000 was annotated using Prodigal version 2.6 [73]. The loci of previously identified *E. coli* O157 sRNA [7] were found in strain 9000 using BLASTn and manually added to strain 9000 .gtf file. Column 3 of the reference GTF file (feature) was manually modified to CDS for all genetic features. Differential expressed (DE) genes were identified with edgeR [74] (p-values =0.05) using the glmQLFit + glmQLFTest parameters. RNA-seq data was uploaded to NCBI Gene Expression Omnibus (GEO) (Accession: **GSE158899**).

### Stx toxin ELISA

3 ml LB was inoculated directly from glycerol stocks and grown overnight at 37 °C. 6 ml LB was inoculated 1/100 from overnight cultures and grown to an OD_600nm_ = 0.6–0.8. Mitomycin C (2 μg/ml) was added and lysis allowed to proceed for 24 h. After 24 h, 1 ml culture was taken and live cells and cell debris removed by centrifugation (13,000 rpm). Stx toxin containing supernatants were further sterilized by syringe filtering (0.22 μm; Milipore). The level of Stx toxin in each sample was assayed using the RIDASCREEN^®^ Verotoxin ELISA kit (R-Biopharm) according to manufacturer guidelines. Differences Stx2 production was assessed by ordinary one-way ANOVA with multiple comparisons where each strain was compared with Z1723.

### Stx Vero cell toxicity

Cytotoxicity of Stx2 toxin was measured on Vero cell monolayers cultured in RPMI medium (Sigma-Aldrich). Cells (100 μl) were plated into 96-well microtitre plates and at ~ 75% confluence the culture medium was replaced with RPMI medium containing diluted (1:1000) Stx2 toxin supernatants. Vero cells were exposed to Stx2 toxin for 72 hours at 37°C, 5% CO_2_. Surviving cells were fixed using paraformaldehyde (2 %) and stained with crystal violet (10 %). Crystal violet was solubilized with 10% acetic acid live/dead cells were quantified spectrophotometrically at 590 nm. Cells exposed to Triton X-100 (0.1 %) and RPMI were used as positive and negative controls for toxicity respectively. Strain toxicity was expressed as a percentage of the toxicity measured for RPMI control. Strain toxicity was analysed by ordinary one-way ANOVA with multiple comparisons where each strain was compared with Z1723.

### Fitness assays

The fitness of strain 9000 variants grown in M9 media was calculated as described previously [75, 76]. Viable-cell counts for each competing strain were determined at time zero (t=0) and again after 24 h of co-culturing by selective plating. Fitness was calculated using the formula:

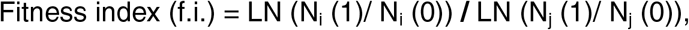

Where N_i_ (0) and N_i_ (1) = initial and final colony counts of strain Z1723 or 9000, respectively and N_j_ (0) and N_j_ (1) = initial and final colony counts of structural variant strain (Z1766, Z1767 or Z1615), respectively

For controls WT strain 9000 or Z1723 were competed against Nalr derivatives generated previously [25]. Fitness was analysed by ordinary one-way ANOVA with Dunnett’s multiple comparisons test where each strain was compared with control.

### RT-qPCR

Total RNA was extracted from cell pellets using a RNeasy® Mini kit (Qiagen) according to manufacturer guidelines. Extracted RNA was quantified and 2 μg of each samples was DNase treated using TURBO DNA-*free*™ kit. 200 ng of DNase treated RNA was then converted to cDNA using iScript™ Reverse Transcription Supermix (Bio-Rad) according to manufacturer guidelines. All qPCR reactions were carried out using iQ™ Syber® Green supermix (Bio-Rad) and *stx2a* specific primers (IDT-DNA): stx2a-F–GAAGAAGATGTTTATGGCGGTTT, stx2a-R–CCCGTCAACCTTCACTGTAA. Cycling conditions were: 95 °C for 15 s (1 cycle), 95 °C for 15 s; 60 °C for 1 min (40 cycles). Gene expression was quantified relative to a standard curve generated from Z1723 genomic DNA.

### Optical mapping

Strains Z1723 and Z1767 were cultured from a single colony in LB medium to an OD_600_ = 0.7. 1 ml of cells/agarose plug were harvested (4000 g, 5 min) and intact chromosomes were extracted according to the Bionano Prep Cell Culture DNA Isolation Protocol (Bionano). Briefly, harvested cells were washed twice in Bionano Cell Buffer (Bionano). Washed cells were embedded in 2 % Low melt agarose plugs and cells were lysed (1 hr at 37°C) with lysozyme enzyme (100 μl) using CHEF Bacterial Genomic DNA Plug Kit (Bio-Rad). DNA containing plugs were washed twice with nuclease free water then treated with Proteinase K (Qiagen) in Bionano Lysis Buffer according to the Bionano Prep Cell Culture DNA Isolation Protocol. All subsequent procedure steps (RNase treatment, DNA extraction, quantitation and labelling) and optical mapping on Bionano Irys platform were provided as a service by Earlham Institute (Norwich, UK). Structural variant analysis was provided by Bionano and structural variant maps visualised using Bionano access (Bionano).

### TTSS expression and secretion

Expression of *ler* and *sepL* was measured using GFP reporter fusion plasmids pDW-LEE1 [77] and pDW6 [78], respectively. Reporter plasmids were transformed into strains Z1723, Z1766 and Z1767 by electroporation and transformants were cultured overnight in LB media supplemented with chloramphenicol (50 μg/ml). Overnight cultures were diluted 1:100 into MEM-HEPES and grown at 37°C (200 rpm) to an OD_600_ 0.8 – 1.0. GFP fluorescence of 200 μL aliquots was measured in a 96-well blank microtiter plate using a FLUOstar Optima plate reader (BMG, Germany). The Gfp promoter-less plasmid pKC26 was used as a control [79].

For EspD secretion, bacteria were cultured in 50 ml of MEM-HEPES at 37°C (200 rpm) to an OD_600_ of 0.8–1.0. Bacterial cells were pelleted by centrifugation at 4000 *g* for 20 min, and supernatants were passed through low protein binding filters (0.45 μm). 10% TCA was used to precipitate proteins overnight, which were separated by centrifugation at 4000 *g* for 30 min at 4°C. The proteins were suspended in 150 μl of 1.5 M Tris (pH 8.8). For bacterial lysates, bacterial pellets were suspended directly in SDS PAGE loading buffer. Proteins were separated by SDS-PAGE using standard methods and Western blotting performed as described previously for EspD and RecA [28].

## Acknowledgements

The authors would like to thank Sandy Fryda-Bradley and the USMARC core sequencing facility for excellent technical assistance. The mention of a trade name, proprietary product, or specific equipment does not constitute a guarantee or warranty by the USDA and does not imply approval to the exclusion of other products that might be suitable. The USDA is an equal opportunity employer and provider.

